# Monitoring mitochondrial translation by pulse SILAC

**DOI:** 10.1101/2021.01.31.428997

**Authors:** Koshi Imami, Matthias Selbach, Yasushi Ishihama

## Abstract

Mitochondrial ribosomes are specialized to translate the 13 membrane proteins encoded in the mitochondrial genome, which shapes the oxidative phosphorylation (OXPHOS) complexes essential for cellular energy metabolism. Despite the importance of mitochondrial translation control, it is challenging to identify and quantify the mitochondrial-encoded proteins due to their hydrophobic nature and low abundance. Here, we introduce a mass spectrometry-based proteomic method that combines biochemical isolation of mitochondria with pulse stable isotope labeling by amino acids in cell culture (pSILAC). Our method provides the highest protein identification rate with the shortest measurement time among currently available methods, enabling us to quantify 12 out of the 13 mitochondrial-encoded proteins. We applied this method to uncover the global picture of (post-)translational regulation of both mitochondrial- and nuclear-encoded subunits of OXPHOS complexes. We found that inhibition of mitochondrial translation led to degradation of orphan nuclear-encoded subunits that are considered to form subcomplexes with the mitochondrial-encoded subunits. The results also allowed us to infer the subcomplex members of each OXPHOS complex. This method should be readily applicable to study mitochondrial translation programs in many contexts, including oxidative stress and mitochondrial disease.

## Introduction

In eukaryotes, both cytosolic and mitochondrial ribosomes (mitoribosomes) play a central role in protein synthesis. Cytosolic ribosomes produce constituents of the cellular proteome encoded in the nuclear genome, while mitoribosomes are specialized to translate the 13 membrane proteins encoded in the mitochondrial genome. These translational products are some of the subunits of oxidative phosphorylation (OXPHOS) complexes that are essential for energy generation in cells. Cytosolic and mitochondrial ribosomes coordinate their translation to enable the proper assembly of OXPHOS complexes on the inner mitochondrial membrane (Couvillion et al., 2016; Dennerlein et al., 2017; Priesnitz and Becker, 2018; Richter-Dennerlein et al., 2016; Tang et al., 2020; Topf et al., 2019). Thus, co-regulation of the mitochondrial and cytosolic translation programs is essential for maintaining mitochondrial proteostasis, i.e., to prevent accumulation of unwanted and potentially harmful assembly intermediates (Isaac et al., 2018). Moreover, many disease-associated mitochondrial mutations are known to impair the mitochondrial translation machinery (Scharfe et al., 2009; Webb et al., 2020), suggesting that dysregulation of mitochondrial translation leads to disease.

Despite the importance of the mitochondrial translation system, a simple and robust method to monitor mitochondrial translation products is lacking. A classical approach is pulse labeling of mitochondrial translation products with radiolabeled amino acids, such as [^35^S]methionine and [^35^S]cysteine (Chomyn, 1996; Lazarou et al., 2007), but the use of radioactive materials and the low resolution of SDS-PAGE gel-based separation of the products limits the utility of this methodology. Alternatively, mass spectrometry (MS)-based proteomic approaches have been developed to monitor protein synthesis (Iwasaki and Ingolia, 2017). Quantitative non-canonical amino acid tagging (QuaNCAT) (Eichelbaum et al., 2012; Howden et al., 2013; Schanzenbächer et al., 2016) relies on pulse labeling of newly synthesized proteins with a methionine analog, azidohomoalanine (AHA) (Dieterich et al., 2006), allowing for selective enrichment of the tagged protein pool through click-chemistry as well as MS-based profiling of the tagged proteins. Nascent chain proteomics using puromycin or its analogs enables isolation and identification of nascent polypeptide chains that are being elongated by the ribosomes (Aviner et al., 2013; Forester et al., 2018; Hünten et al., 2015; Schäfer et al., 2022; Tong et al., 2020; Uchiyama et al., 2020, 2021). However, these methods require a large number of cells (typically >10^7^ cells), involve multiple steps to purify newly synthesized proteins via affinity purification and/or require the isolation of ribosome complexes through density gradient ultracentrifugation.

In contrast, pulse stable isotope labeling of amino acids in cell culture (pSILAC) is a simple and robust technique for global analysis of cellular protein translation (Ebner and Selbach, 2014; Hünten et al., 2015; Imami et al., 2018; Klann et al., 2020; Schwanhäusser et al., 2009; Selbach et al., 2008). pSILAC involves metabolic pulse labeling of newly synthesized proteins with either heavy or medium-heavy amino acids for two cell populations of interest. The newly synthesized (labeled) proteins can be distinguished from pre-existing (non-labeled) proteins by means of MS. The heavy to medium-heavy ratios in the MS spectra reflect the differences in protein production between the two conditions. Of note, a dynamic SILAC approach (Doherty et al., 2009), a variant of SILAC that measures protein turnover, was also recently used to study the turnover rates of mitochondrial proteins in yeast (Saladi et al., 2020) and humans (Bogenhagen and Haley, 2020). Compared to the methods described above, pSILAC does not require many cells (from one to three orders of magnitude fewer) and the downstream experimental process is simply a conventional proteomic workflow. Thus, pSILAC would be a powerful approach to monitor mitochondrial translation.

One of the major challenges in the analysis of mitochondrial translation is that mass spectrometric identification and quantification of the 13 mitochondrial-encoded membrane proteins is hampered by poor protein identification (Figure 1 and see Experimental Procedures) due to the hydrophobic nature and relatively low abundance of these proteins. To overcome this problem, we present a method to comprehensively monitor protein synthesis by mitoribosomes that combines pSILAC with biochemical isolation of mitochondria. Our method offers the highest protein identification rate and the shortest MS measurement time among currently available methods. To demonstrate its utility, we applied it to examine the translational regulation of the mitochondrial- and nuclear-encoded subunits of the OXPHOS complexes.

**Figure 1:**
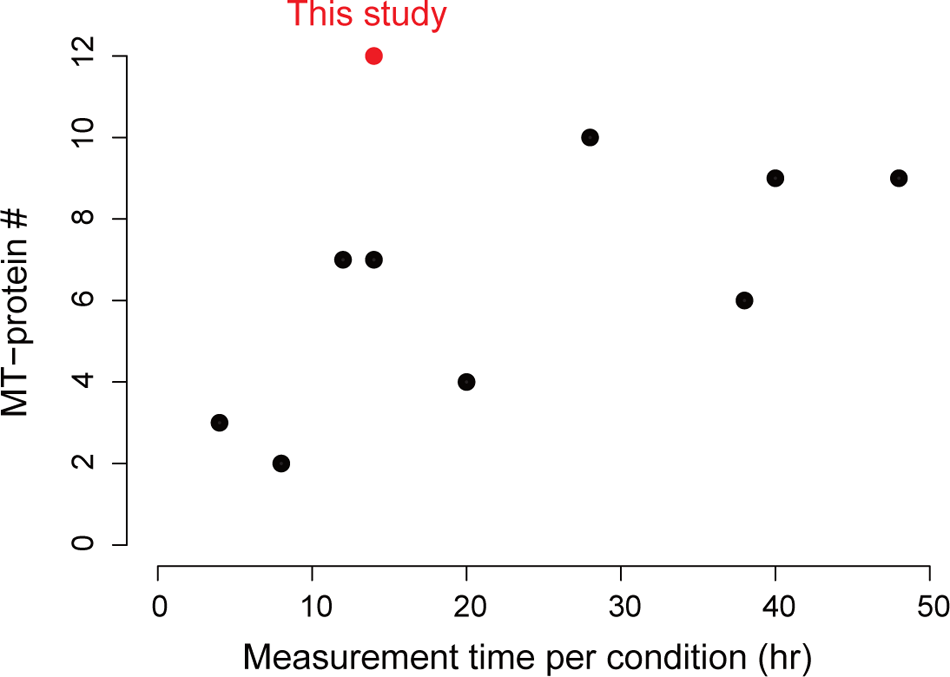
Comparison of the present method with previous studies. The identification numbers of MT-proteins (left axis) and the total LC/MS/MS measurement time (right axis) are shown. The present study gave the highest identification rate of MT-proteins (12 proteins) with the shortest measurement time (14 hr). To compare our method with that of previous reports, we selected studies that had employed pSILAC, AHA, puromycin (and its analog) or dynamic SILAC-tandem-mass tag (TMT). If multiple experiments were performed within a study, the single specific experiment with the highest proteome coverage was chosen. More detailed information is provided in the experimental procedure.

## Results and Discussion

### Biochemical optimization for comprehensive analysis of mitochondrial translation products

While proteomic technologies to quantify protein synthesis have been developed, comprehensive analysis of mitochondrial translation products (hereafter, MT-proteins or MT-encoded subunits) remains challenging, regardless of extensive peptide prefractionation and the use of high sensitivity mass spectrometers (Figure 1 and see Experimental Procedures). To address this issue, we first examined whether the identification of MT-proteins could be improved by combining biochemical isolation of mitochondria with the application of combinations of proteases. Chymotrypsin cleaves on the C-terminal hydrophobic essential amino acids (phenylalanine, tryptophan, and tyrosine) and thus might be suitable for digesting membrane proteins and for pSILAC, which requires essential amino acids for metabolic labeling. We adapted a reported protocol for mitochondrial isolation according to (Frezza et al., 2007), as it is relatively simple (not requiring ultracentrifugation) and quick (approximately 40 min) (see Methods). Mitochondria pellets isolated from HEK293T cells were lysed and protein digestion was performed with 1) chymotrypsin, 2) chymotrypsin-lysC, 3) lysC-trypsin or 4) chymotrypsin-trypsin. In parallel, total cell lysates were digested in the same way as a control. Two biological replicates were analyzed. A complete list of the 6,442 proteins identified is provided in Table S1. We first confirmed that mitochondrial proteins were highly enriched in the isolated mitochondrial fractions, as judged by gene ontology (GO) enrichment analysis (Figure 2A for the top 3 terms and Figure S1 for all terms) and by examination of selected marker proteins (Figure 2B). It should be noted that contamination with proteins from other membranes cannot be completely avoided (Figure S1), as discussed elsewhere (Ma et al., 2017).

**Figure 2:**
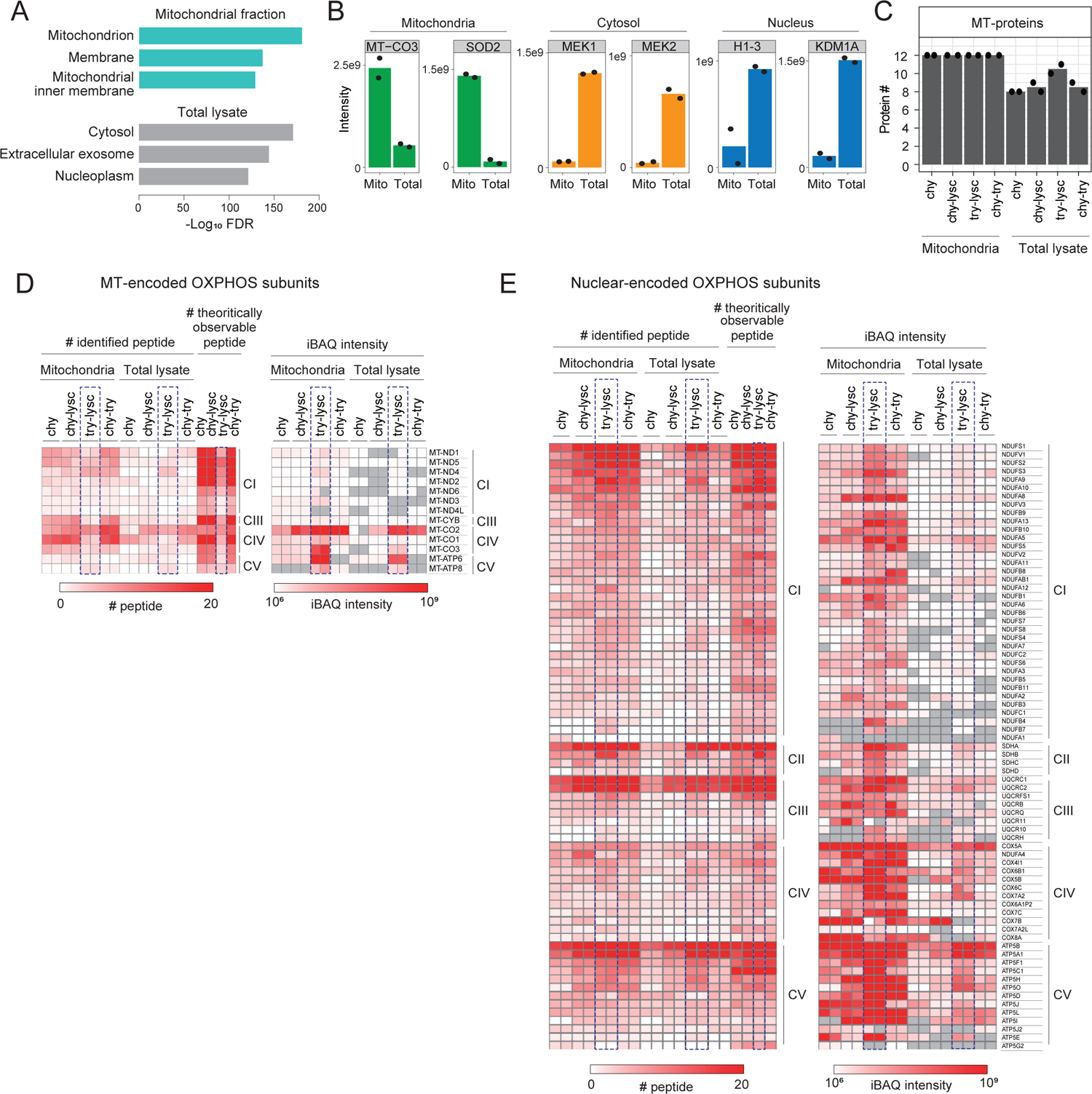
Optimization of biochemical conditions for a proteomic analysis of mitochondrial-encoded proteins. (A) The top3 GO terms enriched in the mitochondrial fraction (top: light green) and total cell lysate (bottom: gray). (B) Abundance profiles of selected organelle (left: mitochondria, center: cytosol, right: nucleus) markers between the mitochondrial fraction (Mito) and total lysate (Total). Only lysC-trypsin digestion samples were shown. (C) The number of identified MT-proteins obtained under 8 different conditions. The bars show the average number of identified proteins from two independent experiments (filled-circle). (D, E) Heatmaps showing the number of identified peptides (left) and iBAQ values (right) from MT- and nuclear-encoded OXPHOS subunits. CI-V, Complex I-V.

Figure 2C shows the number of MT-proteins identified by the different protocols. Isolation of mitochondria significantly enhanced the identification of MT-proteins, as compared to the total cell lysate; In every digestion protocol, 12 of the 13 MT-proteins were identified in the mitochondrial fraction, while on average only 9 proteins were identified in the total lysates. Importantly, it was possible to identify all 13 MT-proteins by utilizing a combination of two digestion protocols (e.g., lysC-trypsin and chymotrypsin) after mitochondrial isolation, indicating that comprehensive profiling of the MT-proteins can be achieved with these two digestion protocols.

To further evaluate the relationship between protease combinations and the number of MT-proteins identified, we assessed theoretically observable (that is, 6<amino acid length<31) and experimentally identified peptides for both MT- and nuclear-encoded OXPHOS subunits (Figures 2D and 2E). We found a clear correlation between the numbers of theoretical and identified peptides for all protease combinations (Figures 2D and 2E left panels). When lysC-trypsin was used, the number of theoretically observable peptides and the number of identified peptides of MT-encoded proteins (blue dish rectangle in Figure 2D left panel) were fewer than those of nuclear-encoded proteins (blue dish rectangles in Figure 2E left panel), indicating that a limited number of peptides with appropriate size is produced from MT-encoded proteins. For other proteases, in contrast, there was no marked difference in the numbers of observable and identified peptides between MT- and nuclear-encoded proteins (Figure 2D and 2E left panels). In addition to the number of identified peptides, we assessed intensity-based absolute quantification (iBAQ) (Schwanhäusser et al., 2011) values (Figures 2D and 2E right panels) which are normalized intensities based on the number of observable peptides. Interestingly, even though fewer peptides were identified from MT-proteins with lysC-trypsin digestion, we observed higher iBAQ intensities for MT-proteins-derived tryptic peptides (Figure 2D and 2E right panels) compared to other protease-cleaved peptides. This observation can potentially be explained by the fact that the physicochemical properties of tryptic peptides with C-terminal positive charges enhance the solubility, ionization efficiency, and/or MS2 fragmentation efficiency of them.

Given that all of the digestion protocols allowed identification of 12 MT-proteins (Figure 2C), we decided to focus on lysC-trypsin digestion of isolated mitochondria for further analysis, as it produced tryptic peptides that are quantifiable using standard pSILAC (Schwanhäusser et al., 2009; Selbach et al., 2008) without modifications to the culture medium, and also provided higher peptide intensities of MT-proteins than did other proteases.

### pSILAC approach to monitor mitochondrial translation

Having established suitable biochemical conditions, we next performed pSILAC experiments. Chloramphenicol (CAP) binds to the A-site crevice on bacterial and mitochondrial ribosomes (Jardetzky, 1963), and thereby inhibits mitochondrial, but not cytosolic translation (Figure 3A). We sought to assess how inhibition of mitochondrial translation through CAP impacts on the synthesis of both MT- and nuclear-encoded OXPHOS complex subunits by applying pSILAC methodology.

**Figure 3:**
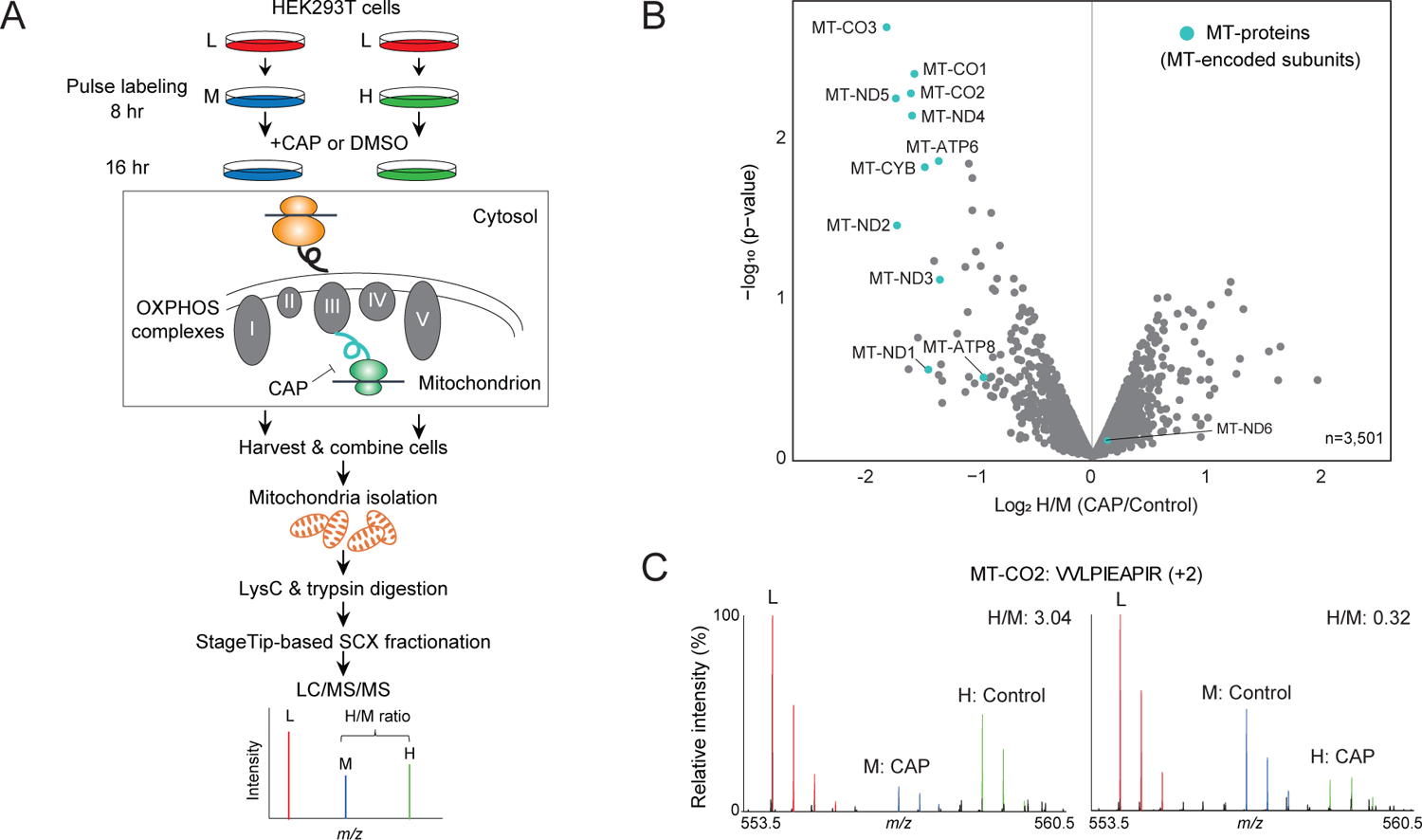
pSILAC experiments. (A) Experimental scheme of pSILAC. HEK293T cells were pulsed-labeled with medium-heavy (M) or heavy (H) amino acids in the presence of CAP or DMSO. Two independent experiments involving a label-swap condition were performed. (B) A volcano plot showing mean log2 fold-change (CAP/DMSO) and -log10 p-value. The MT-proteins are indicated by light green filled circles. (C) MS spectrum of an MT-CO2 peptide (VVLPIEAPIR, +2), as an example.

The experimental scheme of the pSILAC experiment is depicted in Figure 3A. HEK293T cells cultivated in ‘light (L)’ medium were switched to ‘medium-heavy (M)’ or ‘heavy (H)’ medium to pulse-label newly synthesized proteins. Cells were first pre-incubated for 8 hr, and then further incubated for 16 hr in the presence of 10 µg/mL CAP or vehicle (DMSO). We chose 24 hr pulse-labeling, because this period has previously been employed in many studies (Ebner and Selbach, 2014; Hünten et al., 2015; Imami et al., 2018; Klann et al., 2020; Schwanhäusser et al., 2009; Selbach et al., 2008), based on the fact that newly synthesized forms (M and H) of most mammalian proteins can be detected over a time period of 24 hr (Boisvert et al., 2012), thus enabling accurate quantification of H/M ratios. Two independent experiments involving label-swap conditions were performed. The digested peptides were fractionated into 7 fractions using an SCX StageTip (Adachi et al., 2016), and individual fractions were analyzed by means of 110 min LC/MS/MS runs with a 65 min gradient, resulting in a measurement time of approximately 14 hr per condition. We then quantified the H/M ratios in MS spectra to evaluate changes in protein synthesis between CAP and DMSO treatments.

In total, we identified 4,193 proteins, of which 3,501 proteins were quantified in both of two independent experiments and were used for further analysis (see Table S2 for a complete list of proteins and Figure S2 for a more detailed workflow for computational analysis). As expected, our method successfully quantified 12 of the 13 MT-proteins (Figure 3B). Exemplary MS spectra for an MT-CO2-derived peptide (VVLPIEAPIR, +2) are shown in Figure 3C. These results show that the levels of newly synthesized MT-proteins were decreased by CAP treatment (Figures 3B and 3C), confirming that our approach is indeed able to capture the expected changes in mitochondrial translation. Overall, the developed proteomic method provides near-comprehensive (92%) quantification of MT-proteins, representing an improvement of about 2-fold in protein identification as compared with previous pSILAC studies (Figure 1 and see Experimental procedure).

### Relationship between mitochondrial translation and assembly of OXPHOS complexes

In addition to MT-proteins, our mitochondria-focused approach afforded good coverage of OXPHOS complex subunits, including the nuclear-encoded proteins - 41/45 proteins (91%), 4/4 proteins (100%), 9/11 proteins (82%), 14/21 proteins (67%), and 13/17 proteins (76%) from complexes I-V (CI-V), respectively. Therefore, these data can be used to assess how the nuclear-encoded subunits are (post-)translationally regulated in concert with the inhibition of mitochondrial translation. We found significant attenuation of the production of some nuclear-encoded subunits in CI, CIII, and CIV (Figure 4A), though production of most of the nuclear-encoded OXPHOS subunits (especially CII and CV) remained unchanged.

**Figure 4:**
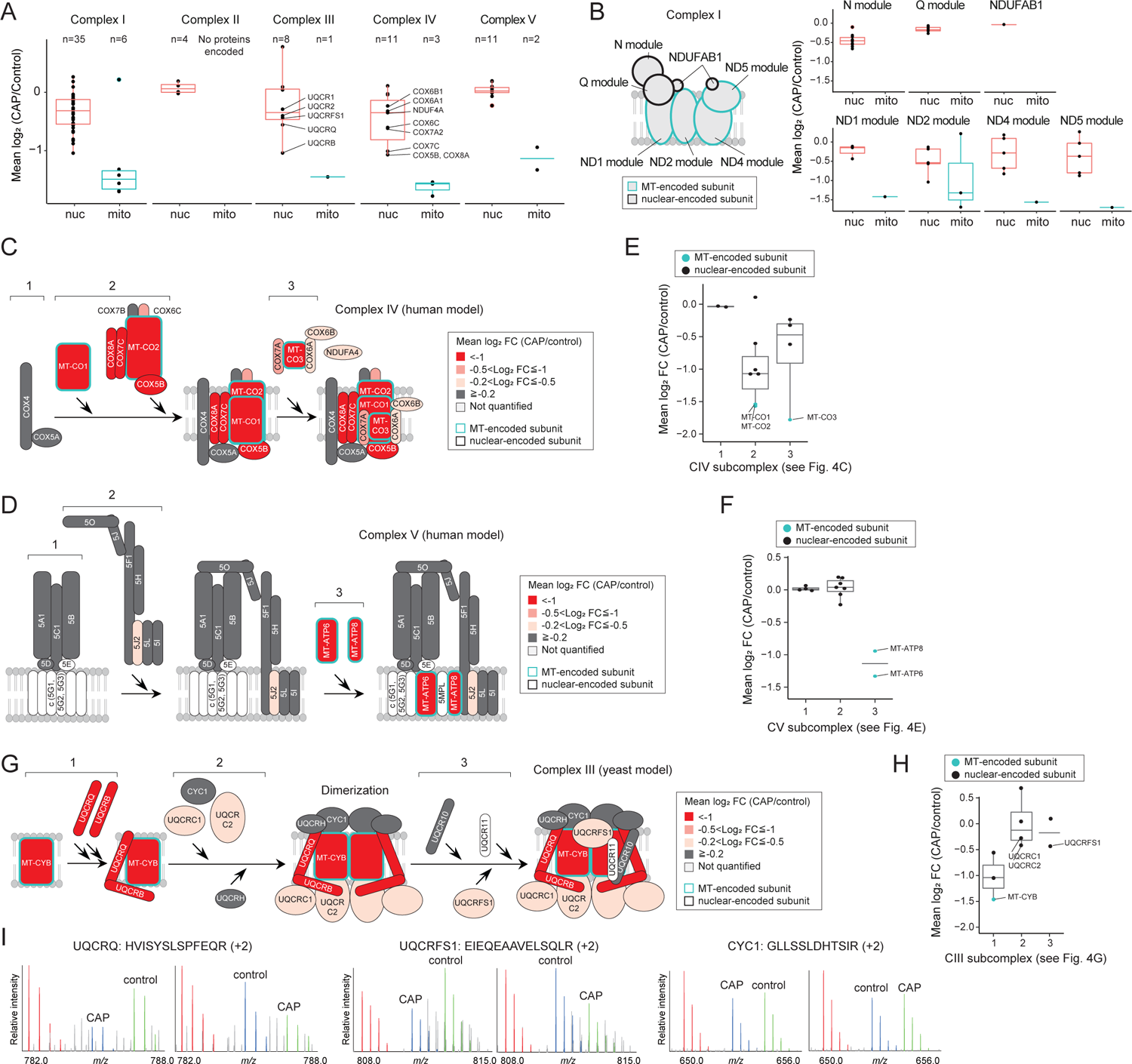
(Post-)translational control and OXPHOS complex assembly. (A) Box plots showing log2 fold-change (CAP/Control) of nuclear-(“nuc”) and mitochondrial-encoded proteins (“mito”) in individual OXPHOS complexes. (B) CI and its subcomplexes. Left: Positions of subunits in complex I (NADH:ubiquinone oxidoreductase) based on humans (Zhu et al., 2016). Right: Boxplots showing log2 fold-change (CAP/Control) for “nuc” and “mito” proteins. (C) An assembly model for complex IV (or cytochrome c oxidase) based on humans (Vidoni et al., 2017). Quantitative values (*i.e.*, log2 FCs (CAP/Control)) are indicated by color. (D) An assembly model for complex V (or ATP synthase) based on humans (He et al., 2018). (E, F) Boxplots showing log2 FCs (CAP/Control) of CIV and CV subunits according to their subcomplexes. (G) A reported assembly model for complex III (cytochrome bc1 complex) based on yeast (Fernández-Vizarra and Zeviani, 2015). (H) Boxplots showing log2 FCs (CAP/Control) of CIII subunits according to its subcomplexes. (I) Exemplary MS spectra for the nuclear-encoded proteins (UQCRQ, CYC1 and UQCRFS1) derived peptides.

To better understand this differential regulation of nuclear-encoded subunits, we focused on CI (NADH:ubiquinone oxidoreductase), CIV (cytochrome c oxidase) and CV (ATP synthase) whose assembly subcomplexes and pathways have been well characterized in humans (Guerrero-Castillo et al., 2017; He et al., 2018; Signes and Fernandez-Vizarra, 2018; Stroud et al., 2016; Vidoni et al., 2017). Overall, we observed a trend for nuclear-encoded subunits that are assembly partners with MT-encoded subunits to be downregulated by CAP (Figure 4). CI is composed of seven modules that are assembled individually (Figure 4B left). Our pSILAC data indicates that some of the nascent nuclear-encoded subunits residing in the same structural modules of the MT-encoded subunits (that is, ND1, ND2, ND4, and ND5-modules) were downregulated (Figure 4B bottom right corner). In contrast, N module, Q module, and NDUFAB1, which are composed of only nuclear-encoded subunits (but not MT-encoded subunits), were rather stable (Figure 4B upper right corner). Likewise, we found that CIV’s nuclear-encoded subunits (COX5B, COX6C, COX7C, and COX8A) affected by CAP are assembly partners of MT-proteins (MT-CO1, MT-CO2, and MT-CO3) (see subcomplex 2 in Figure 4C). In contrast, we observed no significant regulation of the early assembly subunits (COX4 and COX5A) that do not form a subcomplex with MT-proteins (see subcomplex 1 in Figure 4C). Similar results were obtained for CV (Figure 4D); MT-ATP6 and MT-ATP8 are involved in the late step of the complex assembly (step 3 in Figure 4D), and therefore it is reasonable that the early and intermediate subcomplexes of the nuclear-encoded subunits were not regulated (see subcomplexes 1 and 2 in Figure 4D).

These results support the notion that orphan subunits are subject to (post-)translational degradation (Isaac et al., 2018; McShane et al., 2016; Taggart et al., 2020) or a translational pause (Richter-Dennerlein et al., 2016) to prevent accumulation of unwanted assembly intermediates. This is interesting, because it means that our data can throw light on the subcomplex components and/or partners based on quantitative information (that is, H/M ratios of OXPHOS subunits), analogous to genetic studies in which the impacts of the loss of each subunit on the stability of other subunits were investigated (Protasoni et al., 2020; Stroud et al., 2016; Vidoni et al., 2017). To further examine this idea, we grouped the subunits of CIV and CV into three categories based on subcomplexes (see subcomplexes 1-3 in Figures 4C and 4D), and asked whether quantitative information about these subunits can be used to infer the complex components/partners. As expected, we found that H/M (CAP/control) ratios could predict MT-subunit-centric subcomplexes. We observed a trend that nuclear-encoded proteins are co-regulated with MT-proteins within the same subcomplex (Figures 4E and 4F). For example, the nuclear-encoded subunits of CIV in steps 2 and 3 were downregulated (Figure 4E) because their subcomplex assemblies depend on the presence of their partner MT-proteins. On the other hand, the nuclear-encoded subunits of CIV in step 1 and CV in steps 1 and 2 remained unchanged (Figures 4E and 4F) because their subcomplex assemblies were independent of the presence of MT-proteins. These results indicate that subunit members within the same subcomplex module can be inferred based on changes in the levels of newly synthesized subunits.

### Inferring members of intermediate subcomplexes of complex III

We next sought to infer the intermediate steps (subcomplexes) of CIII (cytochrome bc1 complex) assembly, because so far only the first and last steps of its assembly are well understood in humans (Fernández-Vizarra and Zeviani, 2015; Signes and Fernandez-Vizarra, 2018). The assembly model of CIII from yeast (Gruschke et al., 2011, 2012; Hildenbeutel et al., 2014) is shown in Figure 4G. In accordance with the initial step, inhibiting the translation of MT-CYB led to (post-)translational repression of its partner subunits, UQCRQ and UQCRB (see subcomplex 1 in Figures 4G, 4H, and 4I left). Furthermore, consistent with the last step of the assembly in which the Rieske Fe–S protein, UQCRFS1, joins the pre-CIII assembly (Signes and Fernandez-Vizarra, 2018), orphan UQCRFS1 was (post-)translationally downregulated (see subcomplex 3 in Figures 4G and 4H) (see Figure 4I middle for exemplary MS spectra for UQCRFS1). Intriguingly, in contrast to the yeast model, our results imply that UQCRC1 and UQCRC2 may be incorporated into the early-assembled complex of MT-CYB, UQCRQ and UQCRB (see subcomplex 1 in Figures 4G and 4H) during the initial/intermediate steps, because the abundances of newly synthesized UQCRC1 and UQCRC2 decreased in concert with the translational inhibition of MT-CYB (Figures 4G and 4H). In contrast, other subunits (CYC1, UQCRH and UQCR10) appear to be independent of the presence of the early-assembled complex, indicating that these subunits might form distinct module(s) (see Figure 4I right for exemplary MS spectra for CYC1).

### Orphan newly synthesized subunits are more quickly degraded than non-orphan subunits

The data (Figure 4) presented so far indicate that orphan nuclear-encoded OXPHOS subunits whose partner MT-encoded subunits are lost are more likely to be degraded. To confirm this, we performed a slightly modified version of previously reported global pulse-chase experiments (McShane et al., 2016); HEK293T cells were pulse-labeled with H amino acids for 4 hr, followed by chasing newly synthesized H forms for another 4 hr by switching to medium containing M amino acids in the presence of CAP or DMSO (Figure 5A). If newly synthesized (H) proteins are less stable than old (L) proteins their H/L ratios are expected to decrease during the chase. Hence, this experiment should allow us to assess the extent of degradation of newly synthesized proteins during CAP chase by computing the H/L (CAP) / H/L (DMSO) ratios. To this end, we grouped the nuclear-encoded subunits into the two categories; “unchanged (log2 H/M≧-0.5)” or “CAP-sensitive (log2 H/M<-0.5)”, according to the pSILAC experiment (Figures 3 and 4), and asked whether proteins in the CAP-sensitive group (i.e., orphan subunits) are less stable than those in the unchanged group. We first confirmed that CAP inhibited protein synthesis (M-channel) of the CAP-sensitive group more effectively than that of the unchanged group (Figure 5B left panel). Consistent with our hypothesis, we observed a trend that newly made subunits (H-channel) in the CAP-sensitive group were less stable than the other subunits (Figure 5B right panel). We also repeated the pulse-chase experiment and observed the same trend (Figure 5C). Collectively, these results suggest that protein degradation is a key pathway of elimination of orphan nuclear-encoded subunits that cannot form a subcomplex with MT-encoded subunits.

**Figure 5:**
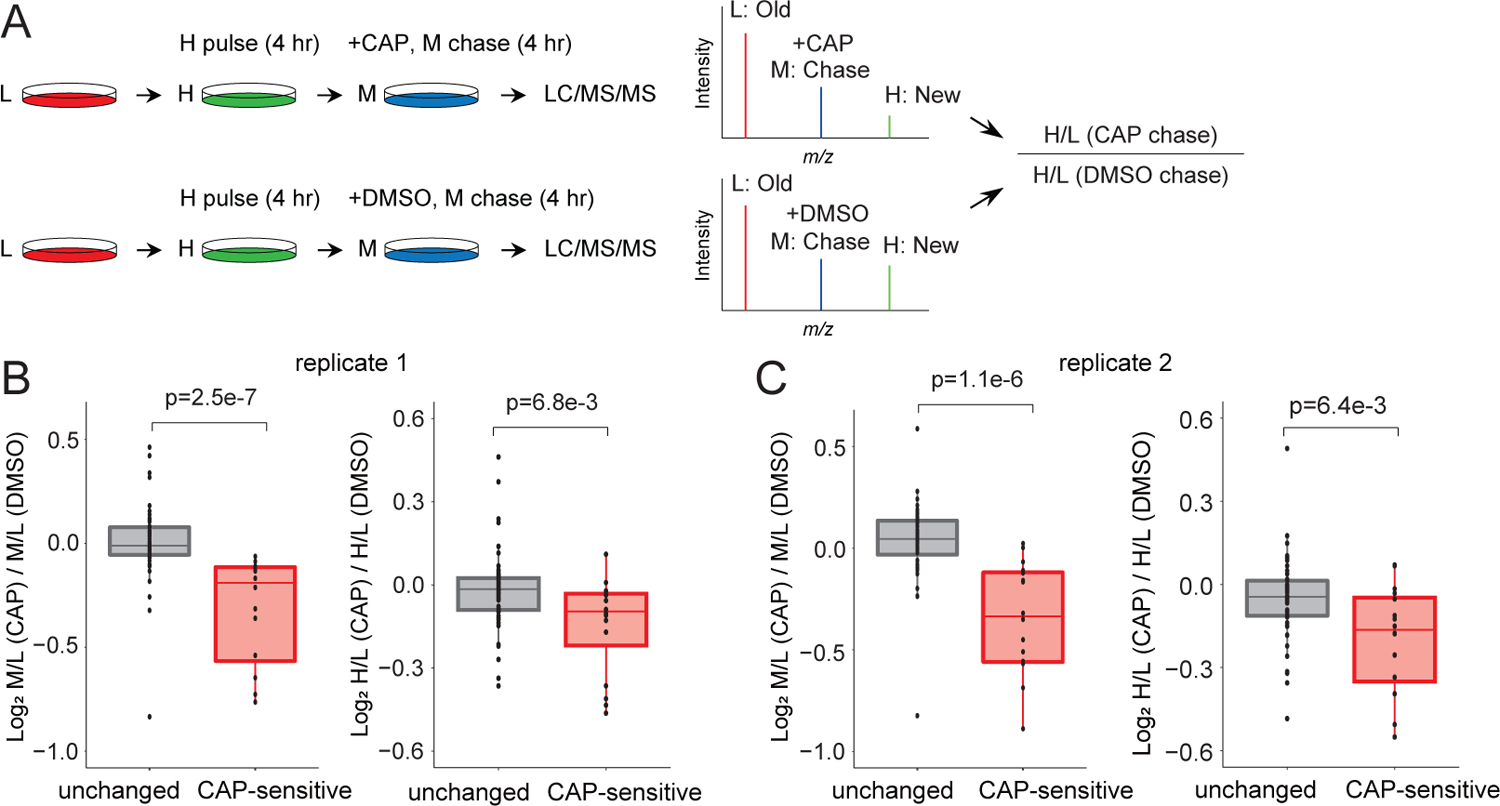
Orphan nuclear-encoded OXPHOS subunits are degraded after protein synthesis. (A) Experimental scheme of a global pulse-chase experiment. HEK293T cells were pulse-labeled with H amino acids for 4 hr, followed by chasing newly synthesized H forms for 4 hr by switching to medium containing M amino acid in the presence of CAP or DMSO. The degree of degradation of newly synthesized H proteins can be assessed based on H/L (CAP) / H/L (DMSO). (B) Effects of CAP on protein synthesis and degradation of nuclear-encoded subunits. A boxplot showing the degree of inhibition of protein synthesis of newly made M forms between CAP and DMSO treatments (left panel). A box plot showing the degree to which newly made H forms are degraded by CAP (right panel). “Unchanged” and “CAP-sensitive” represent nuclear-encoded subunits whose protein synthesis was unchanged and inhibited by CAP, respectively (see Figure 4). P-value was computed using the two-sided Wilcoxon rank-sum test. (C) Same as in (B) but the result was from another independent experiment. P-value was computed using the two-sided Wilcoxon rank-sum test.

## Conclusions

While pSILAC is an established approach for studying protein synthesis, it has never been applied to study mitochondrial translation. The significance of this study lies in the improvement and development of the pSILAC approach combined with a simple biochemical separation for mitochondria. To our knowledge, this is the first study to achieve a near-comprehensive profiling of nascent MT-proteins translated by mitoribosomes. Moreover, this methodology provides a global view of OXPHOS complex assembly on the basis of (post-)translational regulation of mitochondrial- and nuclear-encoded proteins. We found that CAP-mediated inhibition of mitochondrial translation induced degradation of the nuclear-encoded proteins of OXPHOS complexes, and this regulation appears to be maintained at the structural module level; our results suggest that orphan nascent nuclear-encoded proteins are degraded in concert with the loss of their partner MT-proteins in the same structural module. We believe that this methodology will enable us to probe mitochondrial translation programs in many contexts, including oxidative stress and mitochondrial disease.

## Acknowledgements

We are very grateful to Erik McShane (Harvard Medical School) for critical reading of the manuscript. We thank the members of the Department of Molecular & Cellular BioAnalysis and the Department of Proteomics and Drug Discovery for fruitful discussion. KI thanks the Samuro Kakiuchi Memorial Research Award for Young Scientists for supporting this study. This work was supported by JSPS KAKENHI Grant-in-Aid for Scientific Research (Grant Numbers 18K14674, 20H03241, 20H04844, 21H05720 to KI and 17H05667 to YI), the Takeda Science Foundation to KI, JST Strategic Basic Research Program CREST (18070870), and AMED Advanced Research and Development Programs for Medical Innovation CREST (18068699) to YI.

## Author contributions

Conceptualization & Methodology, K.I.; Investigation, K.I and Y.I.; Formal Analysis, K.I., M.S. and Y.I; Resources, K.I, M.S., and Y.I; Writing - Original Draft, K.I; Writing - Review & Editing, M.S. and Y.I., Visualization, K.I; Supervision, K.I. and Y.I.; Funding Acquisition, K.I., M.S., and Y.I.

## Supplemental Information

Table S1: A list of proteins identified in mitochondrial fraction and total lysate (related to Figure 2)

Table S2: A list of proteins quantified in pSILAC experiments (related to Figures 3 and 4)

Table S3: A list of OXPHOS subunits quantified in global pulse-chase experiments (related to Figure 5)

## Experimental procedures

### Cell culture and pulse labeling

HEK293T cells obtained from American Type Culture Collection were cultured in Dulbecco’s modified Eagle’s medium (DMEM) (Fujifilm Wako, Osaka, Japan) containing 10% fetal bovine serum (FBS) (Thermo Fisher Scientific, Waltham, USA) in 10 cm diameter dishes. All cells were maintained in a humidified 37°C incubator with 5% CO_2_. For pulse SILAC labeling (related to Figure 3), the cell culture medium was switched to arginine- and lysine-free DMEM (Thermo Fisher Scientific, Waltham, USA) supplemented with 10% FBS and either “heavy” amino acids [0.398 mM L-(^13^C_6_,^15^N_4_)-arginine (Arg”10”) and 0.798 mM L-(^13^C_6_,^15^N_2_)-lysine (Lys”8”)] or “medium-heavy” amino acids [0.398 mM L-(^13^C_6_)-arginine (Arg”6”) and 0.798 mM L-(D_4_)-lysine (Lys”4”)] (Cambridge Isotope Laboratories, Tewksbury, USA). For chloramphenicol (CAP) (Fujifilm Wako) treatment, HEK293T cells were first pre-incubated with the corresponding SILAC medium for 8 hr, and then further incubated for 16 hr in the presence of 10 µg/mL CAP or vehicle (DMSO). The cells were washed and harvested in ice-cold PBS, and pelleted by centrifugation at 600 × *g* for 5 min at 4°C. Label-swap (biological duplicate) experiments were performed. For global pulse-chase experiments (related to Figure 5), cells were pulse-labeled with “heavy” amino acids for 4 hr as described above, followed by chasing newly synthesized “heavy” forms for 4 hr by switching to medium containing “medium-heavy” amino acids in the presence of 10 µg/mL CAP. Two independent experiments of the pulse-chase experiments under the CAP condition were performed. As a vehicle control, cells were chased in the presence of 0.1% DMSO instead of CAP and used as a universal reference.

### Mitochondria isolation

Mitochondria were isolated as reported (Frezza et al., 2007). For pSILAC samples, corresponding medium-heavy and heavy-labeled cells were combined at this stage. The cell pellets were resuspended in 500 µL of ice-cold mitochondria isolation buffer (10 mM Tris-MOPS [3-(N-morpholino)propanesulfonic acid] pH 7.4 containing 1 mM EGTA/Tris and 200 mM sucrose). The cells were homogenized using a glass/Teflon Potter Elvehjem homogenizer (2 mL volume; 40 strokes). The homogenate was transferred to a new 1.5 mL tube and centrifuged at 600 *x g* for 10 min at 4°C. The supernatant was transferred to a new 1.5 mL tube and centrifuged at 7,000 *x g* for 10 min at 4°C. The pellet containing mitochondria was resuspended in 100 µL of ice-cold mitochondria isolation buffer and centrifuged at 7,000 *x g* for 10 min at 4°C. The supernatant was discarded, and the pellet was used as the mitochondrial fraction.

### Protein digestion

Protein digestion was performed according to the phase-transfer surfactant (PTS)-aided digestion protocol, as described previously (Masuda et al., 2008, 2009). Briefly, the mitochondrial fraction was lysed with PTS buffer [12 mM sodium deoxycholate (SDC) (Fujifilm Wako), 12 mM sodium *N*-lauroyl sarcosinate (SLS) (Fujifilm Wako) in 0.1 M Tris-HCl pH 8.0] and incubated with 10 mM dithiothreitol (DTT) at 37°C for 30 min, followed by alkylation with 50 mM iodoacetamide at 37°C for 30 min in the dark. The samples were diluted 5 times with 50 mM ammonium bicarbonate. To optimize the digestion protocol for mitochondrial translation products, proteins were digested with 1) chymotrypsin (Promega, Madison, USA), 2) chymotrypsin and lysyl endopeptidase (lys-C) (Fujifilm Wako) 3) lys-C and trypsin (Promega) and 4) chymotrypsin and trypsin at a protein-to-protease ratio of 50:1 (w/w) overnight at 37°C on a shaking incubator. For the pSILAC experiments, proteins were first digested with lys-C for 3 hr at 37°C and then with trypsin overnight at 37°C on a shaking incubator. Next day, an equal volume of ethyl acetate (Fujifilm Wako) was added to the sample and digestion was quenched by adding 0.5% trifluoroacetic acid (TFA) (final concentration). The samples were shaken for 1 min and centrifuged at 16,000 *x g* for 2 min at 25°C. The organic phase containing SDC and SLS was discarded. The resulting peptide solution was evaporated in a SpeedVac and the residue was resuspended in 200 µL 0.1% TFA 5% acetonitrile (ACN). The peptides were desalted with an SDB-XC StageTip (Rappsilber et al., 2003) or fractionated into 7 fractions using an SDB-XC-SCX StageTip (Adachi et al., 2016). Each sample solution was evaporated in a SpeedVac and the residue was resuspended in 0.5% TFA and 4% ACN.

### LC/MS/MS analysis

Nano-scale reversed-phase liquid chromatography coupled with tandem mass spectrometry (nanoLC/MS/MS) was performed on an Orbitrap Fusion Lumos mass spectrometer (Thermo Fisher Scientific), connected to a Thermo Ultimate 3000 RSLCnano pump and an HTC-PAL autosampler (CTC Analytics, Zwingen, Switzerland) equipped with a self-pulled analytical column (150 mm length × 100 μm i.d.) (Ishihama et al., 2002) packed with ReproSil-Pur C18-AQ materials (3 μm, Dr. Maisch GmbH, Ammerbuch, Germany). The mobile phases consisted of (A) 0.5% acetic acid and (B) 0.5% acetic acid and 80% ACN. For pSILAC experiments, peptides were eluted from the analytical column at a flow rate of 500 nL/min by altering the gradient: 5-10% B in 5 min, 10-40% B in 60 min, 40-99% B in 5 min and 99% for 5 min, and a 300 min gradient was used for the biochemical optimization. The Orbitrap Fusion Lumos instrument was operated in the data-dependent mode with a full scan in the Orbitrap followed by MS/MS scans for 3 sec using higher-energy collisional dissociation (HCD). The applied voltage for ionization was 2.4 kV. The full scans were performed with a resolution of 120,000, a target value of 4×10^5^ ions, and a maximum injection time of 50 ms. The MS scan range was *m/z* 300–1,500. The MS/MS scans were performed with 15,000 resolution, 5×10^4^ target value, and 50 ms maximum injection time. The isolation window was set to 1.6, and the normalized HCD collision energy was 30. Dynamic exclusion was applied for 20 sec.

### Database searching and protein quantification

All raw files were analyzed and processed by MaxQuant (v1.6.0.13 or 1.6.15.0) (Cox and Mann, 2008). Search parameters included two missed cleavage sites and variable modifications such as methionine oxidation, protein N-terminal acetylation, and SILAC-specific modifications [L-(^13^C_6_,^15^N_4_)-arginine, L-(^13^C_6_,^15^N_2_)-lysine, L-(^13^C_6_)-arginine, L-(D_4_)-lysine]. Cysteine carbamidomethylation was set as a fixed modification. The peptide mass tolerance was 4.5 ppm, and the MS/MS tolerance was 20 ppm. The database search was performed with Andromeda (Cox et al., 2011) against the UniProt/Swiss-Prot human database (downloaded on 2014-10) with common serum contaminants and enzyme sequences. The false discovery rate (FDR) was set to 1% at the peptide spectrum match (PSM) level and protein level. The ‘match between runs’ functions were employed. For protein quantification, a minimum of one unique peptide ion was used, and to ensure accurate quantification, we required proteins to be quantified in all samples for further analysis. Protein intensities from SILAC medium-heavy and heavy channels were normalized using the variance stabilization normalization (Huber et al., 2002) in the R package of DEP (Zhang et al., 2018) to correct for mixing error between the two SILAC labeled lysates. P-values were computed based on differential expression of proteins using protein-wise linear models and empirical Bayes statistics through the *limma* function (Ritchie et al., 2015).

### Assessment of the number of identified mitochondrial translation products and measurement time (related to Figure 1)

There are a number of studies on cellular translation using pSILAC, AHA and/or puromycin, as described in the introduction. To compare our findings with those of previous reports, we chose studies that had employed pSILAC with medium-heavy and heavy amino acids (*i.e*., triplex SILAC) and that showed reasonably high proteome coverage (see Table 1). Recent studies using AHA, puromycin (and its analog) or dynamic SILAC-tandem-mass tag (TMT) were also included. If multiple experiments were performed within a study, the single specific experiment with the highest proteome coverage was chosen (see Figure 1). The measurement time per experiment is a conservative estimate, as some studies provided only the LC gradient time and did not mention total measurement time.

**Table 1:** Method comparison

| Reference | This study | Ebner et al 2014 | Hunten et al 2015 | Imami et al 2018 | Klann et al 2020 | Schanzenbächer et al 2016 | Aviner et al 2013 | Tong et al 2020 | Uchiyama et al 2020 | Adriana Schaefer et al 2021 |
| --- | --- | --- | --- | --- | --- | --- | --- | --- | --- | --- |
| PubMed ID | - | 24637697 | 26183718 | 30220558 | 31812349 | 27764671 | 23934657 | 32531160 | 32926143 | 34847359 |
| Measurement time (h) | 14 | 38 | 20 | 8 | 48 | 12 | 4 | 40 | 14 | 28 |
| #MT-proteins | 12 | 6 | 4 | 2 | 9 | 7 | 3 | 9 | 7 | 10 |
| # total proteins | 3501 | 5435 | 5126 | 3422 | 6039 | 5940 | 2535 | 2589 | 4365 | 4074 |
| Methodology | pSILAC | pSILAC | pSILAC | pSILAC | pSILAC-TMT | AHA | Puro | Puro-TMT | Puro-pSILAC | TMT-pSILAC |
| Cell type | HEK293T | HeLa | SW480 | HEK293T | HeLa | Mouse neurons | HeLa | THP-1 | HeLa | HeLa |
| MS | Fusion Lumos | LTQ-Orbitrap | LTQ-Orbitrap Elite | Q-Exactive Plus | Fusion Lumos | Q-Exactive Plus | Q-Exactive | LTQ-Orbitrap Elite | Fusion Lumos | Fusion Lumos |
| Fractionation | StageTip-based SCX fractionation (7 fractions) | GeLCMS (15 slices) | GeLCMS (20 slices) | Single shot | High pH-reversed phase fractionation (24 fractions) | Single shots (4 times) | Single shot | High pH-reversed phase fractionation (20 fractions) | StageTip-based SCX fractionation (7 fractions) | High pH-reversed phase fractionation (8 fractions) |

### Gene ontology (GO) enrichment analysis (related to Figure 2A)

GO enrichment analysis was performed using DAVID (Huang et al., 2009). To identify enriched GO terms in the mitochondrial fraction and total cell lysate, we used only proteins quantified in all of the digestion protocols. The top 3 enriched terms for cellular components are shown in Figure 2A, while a full list of enriched GO terms is shown in Figure S1. FDRs were corrected by the Benjamini-Hochberg method.

### Analysis of global pulse-chase experiment (related to Figures 5D and 5E)

Only nuclear-encoded OXPHOS subunits were analyzed, and they were grouped into the two categories, “unchanged (log2 H/M≧-0.5)” or “CAP-sensitive (log2 H/M<-0.5)”, based on the pSILAC experiment shown in Figures 3 and 4. Heavy (H) peaks in MS spectra represent the abundance of newly synthesized proteins after the chase in the presence of CAP or DMSO. Light (L) peaks indicate pre-existing proteins. Hence, the degree of degradation of newly synthesized H proteins induced by CAP treatment can be assessed by computing H/L (CAP) / H/L (DMSO). Individual dots in the box plots (Figures 5D and 5E) represent log2 [H/L (CAP) / H/L (DMSO)] values for individual nuclear-encoded subunits categorized into the “unchanged” or “CAP-sensitive” group. The p-values were computed using the two-sided Wilcoxon rank-sum test.

### Data and software availability

The proteomics data have been deposited to the ProteomeXchange Consortium via jPOST (Moriya et al., 2019; Okuda et al., 2017) partner repository with the dataset identifier JPST001007 (PXD022476 for ProteomeXchange). URL https://repository.jpostdb.org/preview/1782746175fac8dd943bc0

### Quantification and statistical analysis

The type of statistical test (e.g., t-test using protein-wise linear models and empirical Bayes statistics) is annotated in the Figure legend and/or in the Methods and Resources segment specific to the analysis. In addition, statistical parameters such as the value of n, mean/median and significance level are reported in the Figures and/or in the Figure Legends. Statistical analyses were performed using R as described in Methods and Resources for each individual analysis.

---

**Figure S1:**
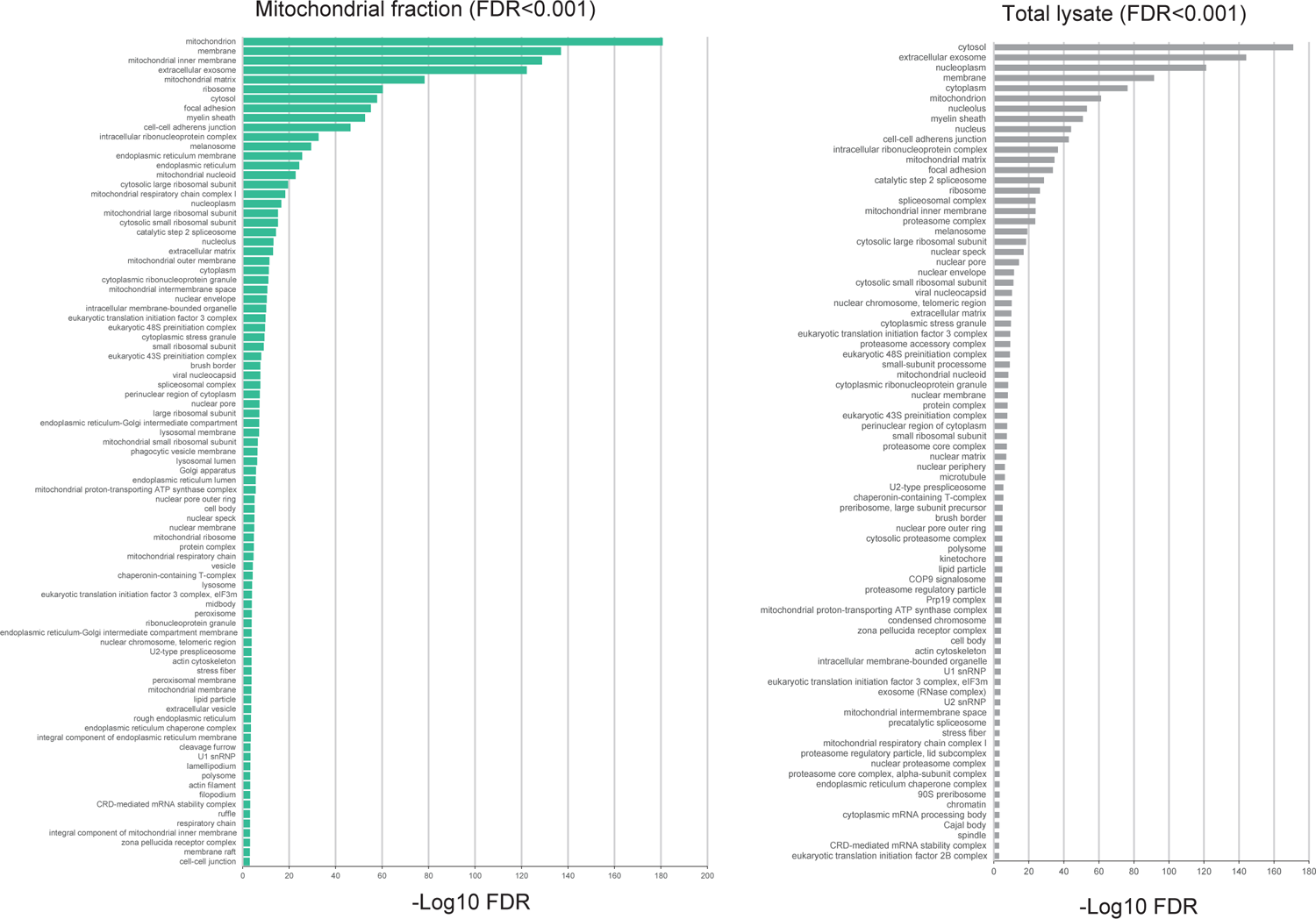
Enriched GO terms in mitochondrial fraction-(left) and total lysate-(right) derived samples, related to Figure 2A.

**Figure S2:**
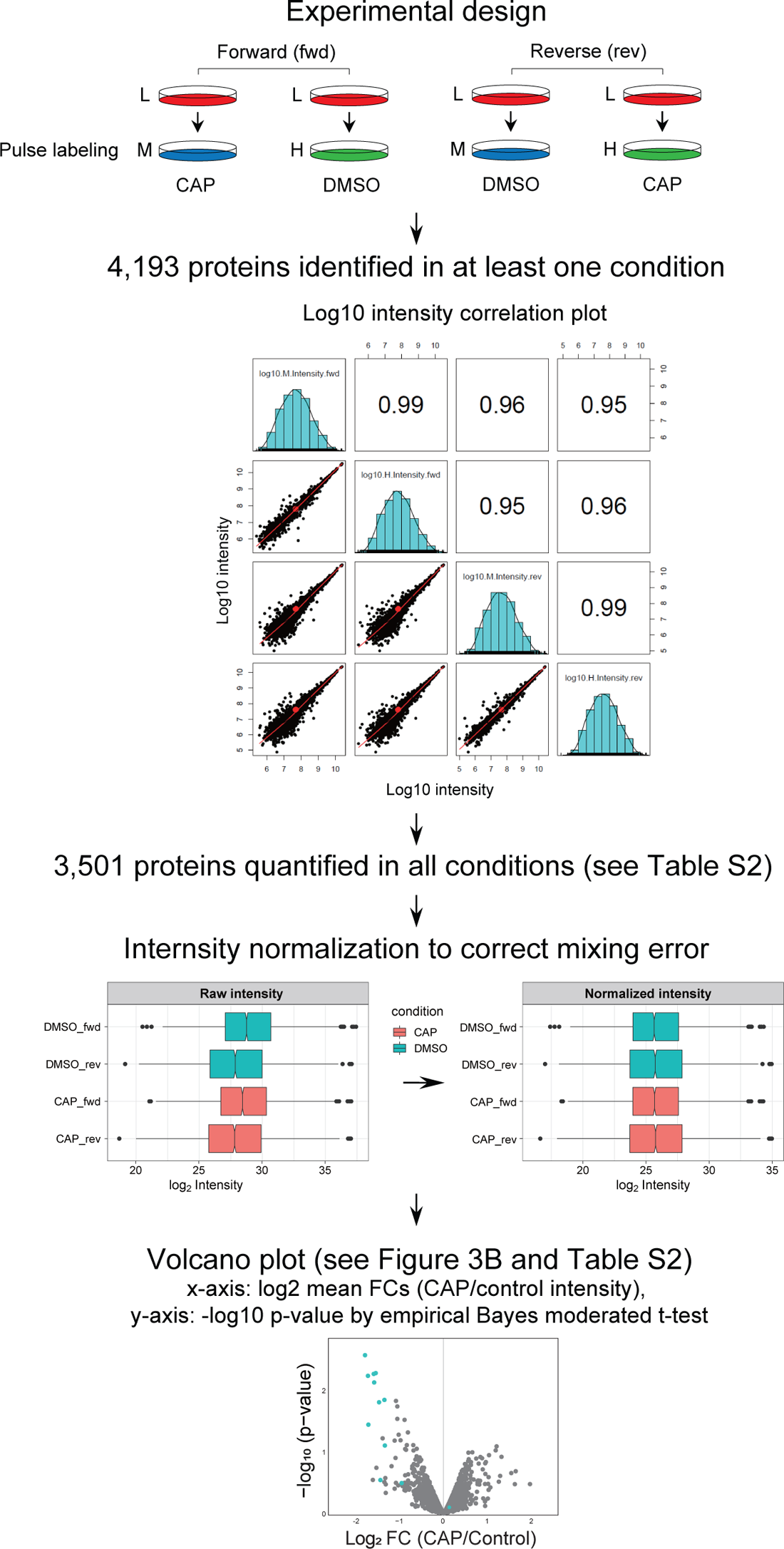
A workflow for the analysis of the pSILAC data related to Figures 3 and 4. Two independent pSILAC experiments (forward and reverse label-swap) were performed; of the 4,193 proteins identified under at least one condition, 3,501 proteins quantified under all conditions were used for the following analysis. To correct for the mixing ratio of H and M samples, variance stabilization normalization (Huber et al., 2002) was performed. Finally, mean log2FC and t-test were used to obtain a volcano plot (Figure 3B)

